# The potato NLR immune receptor R3a does not contain non-canonical integrated domains

**DOI:** 10.1101/056242

**Authors:** Artemis Giannakopoulou, Angela Chaparro-Garcia, Sophien Kamoun

## Abstract

A recent study by Kroj et al. (New Phytologist, 2016) surveyed nucleotide binding-leucine rich repeat (NLR) proteins from plant genomes for the presence of extraneous integrated domains that may serve as decoys or sensors for pathogen effectors. They reported that a FAM75 domain of unknown function occurs near the C-terminus of the potato late blight NLR protein R3a. Here, we investigated in detail the domain architecture of the R3a protein, its potato paralog R3b, and their tomato ortholog I2. We conclude that the R3a, R3b, and I2 proteins do not carry additional domains besides the classic NLR modules, and that the FAM75 domain match is likely a false positive among computationally predicted NLR-integrated domains.

---

Plants use immune receptors to recognize invading pathogens and pests. The largest family of intracellular immune receptors is the nucleotide binding-leucine rich repeat (NB-LRR or NLR) protein family - an important element of defense against pathogens in both plants and animals (Maekawa et al. 2011, Jacob et al. 2013). NLR proteins recognize pathogen-secreted molecules, termed effectors, which translocate into the host cytoplasm, triggering an immune response that leads to resistance (Jones and Dangl 2006, Ooijen et al. 2007, Dodds and Rathjen 2010, Win et al. 2012). NLR recognition of pathogen effectors can be direct through physical interaction between receptor and effector or indirect via receptor monitoring of effector-mediated modifications of plant target proteins (Mackey et al. 2002, Axtell and Staskawicz 2003, Rooney et al. 2005). Another model of NLR-detection of pathogen effectors has recently been described in which an extraneous domain, derived from a host target, has integrated into the NLR protein to enable effector recognition (Cesari et al. 2014, Nishimura et al. 2015, Wu et al. 2015). These “integrated domains” have been described as decoys, sensors, and baits, and appear diverse and widespread among plant NLR proteins (Cesari et al. 2014, Nishimura et al. 2015, Wu et al. 2015, Khan et al. 2016, Kroj et al. 2016, Sarris et al. 2016). To determine the extent to which NLR proteins carry integrated domains, Kroj et al (2016), surveyed NLRs from the genome sequences of 31 plant species for non-canonical domains. Among their predictions, they reported that a FAM75 domain of unknown function occurs near the C-terminus of the potato late blight resistance protein R3a (Kroj et al. 2016).

R3a is an NLR protein that confers resistance to the Irish potato famine pathogen *Phytophthora infestans* (Huang et al. 2005). It is one of the first *R* genes to be used in potato breeding for resistance against the late blight pathogen *Phytophthora infestans* (Armstrong et al. 2005, Bos et al. 2006, Morgan and Kamoun 2007, Birch et al. 2008). *R3a* carries a predicted coiled-coil (CC) domain in its N-terminal part (Huang et al. 2005) and shares 88% nucleotide identity and 83% amino acid similarity with *I2,* a tomato NLR that mediates resistance to the soil-borne fungus *Fusarium oxysporum f. sp. lycopersici* (Ori et al. 1997, Simons et al. 1998). The R3 locus includes another functional NLR gene, *R3b*, which shares 82% nucleotide identity with R3a (Huang et al. 2004, Li et al. 2011). R3b also confers resistance to *P. infestans* but with a distinct recognition specificity compared to R3a (Huang et al. 2004, Li et al. 2011). R3a and R3b share their highest similarity in the CC domain (79% amino acid identity), whereas R3a and I2 are mostly similar in the nucleotide binding (NB) domain (86% amino acid identity) (Huang et al. 2005, Li et al. 2011).

In their survey of NLR-integrated domains, Kroj et al. (2016), reported that R3a contains a non-canonical domain, termed FAM75 (see Figure 1 of Kroj et al. 2016). This report surprised us given that R3a and other members of the R3/I2 superfamily are fairly compact NLRs with little room for extraneous domains in addition to the classic modules. Here, we revisited the domain architecture of R3a to check the robustness of this finding.

## RESULTS AND DISCUSSION

To determine whether R3a carries an integrated domain in addition to classic NLR modules, we reexamined an alignment between R3a, its paralog R3b, and their tomato ortholog I2 (Figure 1). The three proteins diverge mainly in the leucine rich repeat (LRR) domain, yet all of them share 29 xxLxLxx repeat units (Figure 1A). Notably, the R3a region between amino acids (aa) 1148-1263, predicted to encode the FAM75 domain, clearly overlaps with five LRRs (Figure 1B).

**Figure 1.**
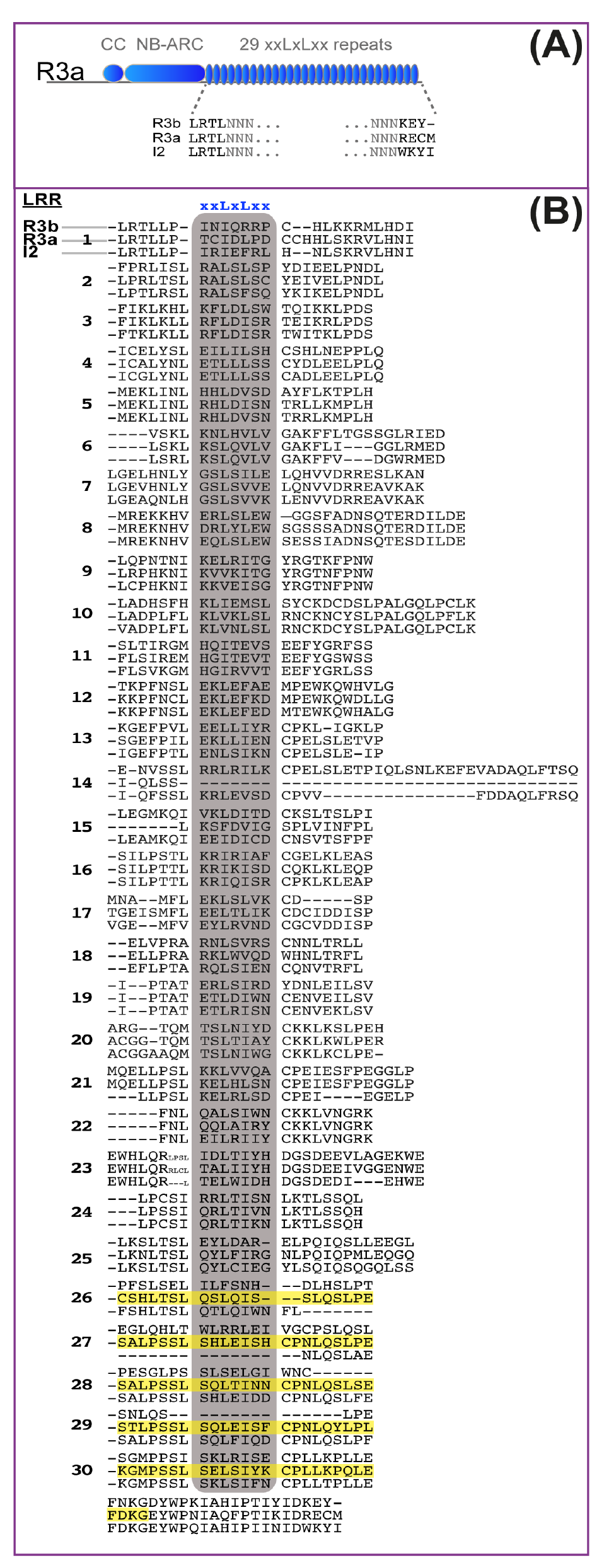
R3a does not carry a FAM75 domain. A, Schematic diagram of the R3aprotein. The LRR domains of R3b, R3a and I2 are shown. B, Alignment of the LRR domains of R3b (top), R3a (middle) and I2 (bottom). The LRR units in the alignment are numbered. The consensus xxLxLxx of each repeat unit is highlighted with a gray box. R3a aa residues 1148-1263 predicted by Kroj et al. (2016) to encode a FAM75 domain are highlighted in yellow. The following protein sequences were used: R3b (GenBank accession AEC47890.1) R3a (GenBank accession AAW48299.1) and I2 (GenBank accession ALF36835.1).

To further examine the domain architecture of R3a, we performed an InterPro protein sequence search (Mitchell et al. 2015). This failed to identify additional domains besides the LRR in the C-terminal half of R3a, consistent with the alignment and manual annotation of the LRR (Figure 2).

**Figure 2.**
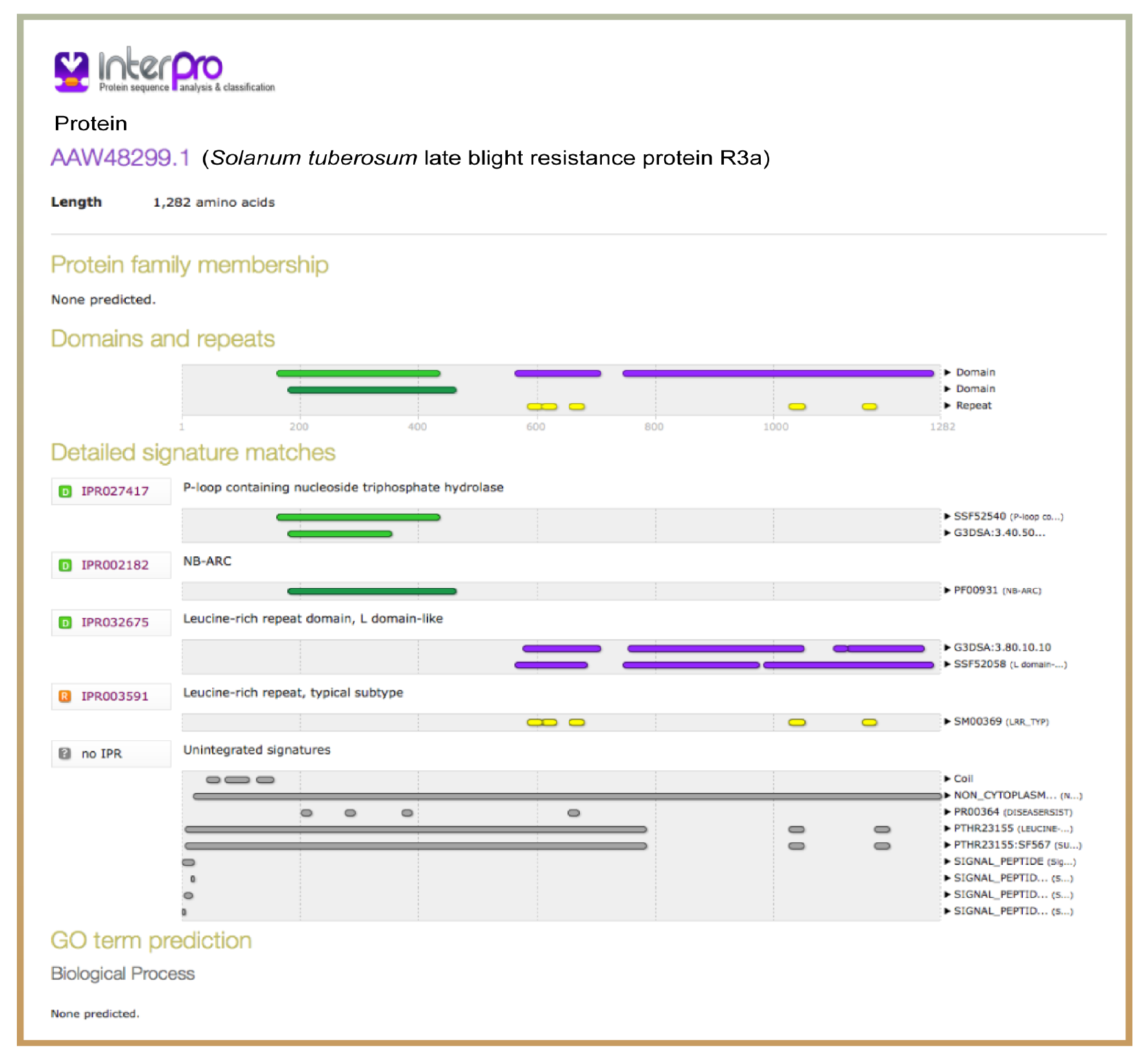
R3a protein domain architecture. We applied the InterPro protein sequence tool to the R3a sequence (Mitchell et al. 2015). Predicted domains and important sites of R3a are shown. The following R3a protein sequence was used for the analysis (GenBank accession AAW48299.1).

Next, we performed homology structure modeling of the R3a LRR region based on protein fold recognition algorithms implemented by IntFold (Figure 3) (Buenavista et al. 2012). We modeled the R3a LRR domain using as a template the top scoring Intfold protein, the Toll-like receptor 13 (PDB code 4z0c) (Song et al. 2015). The homology model confirmed that the presumed FAM75 region overlaps with the last five LRRs of R3a (Figure 3).

**Figure 3.**
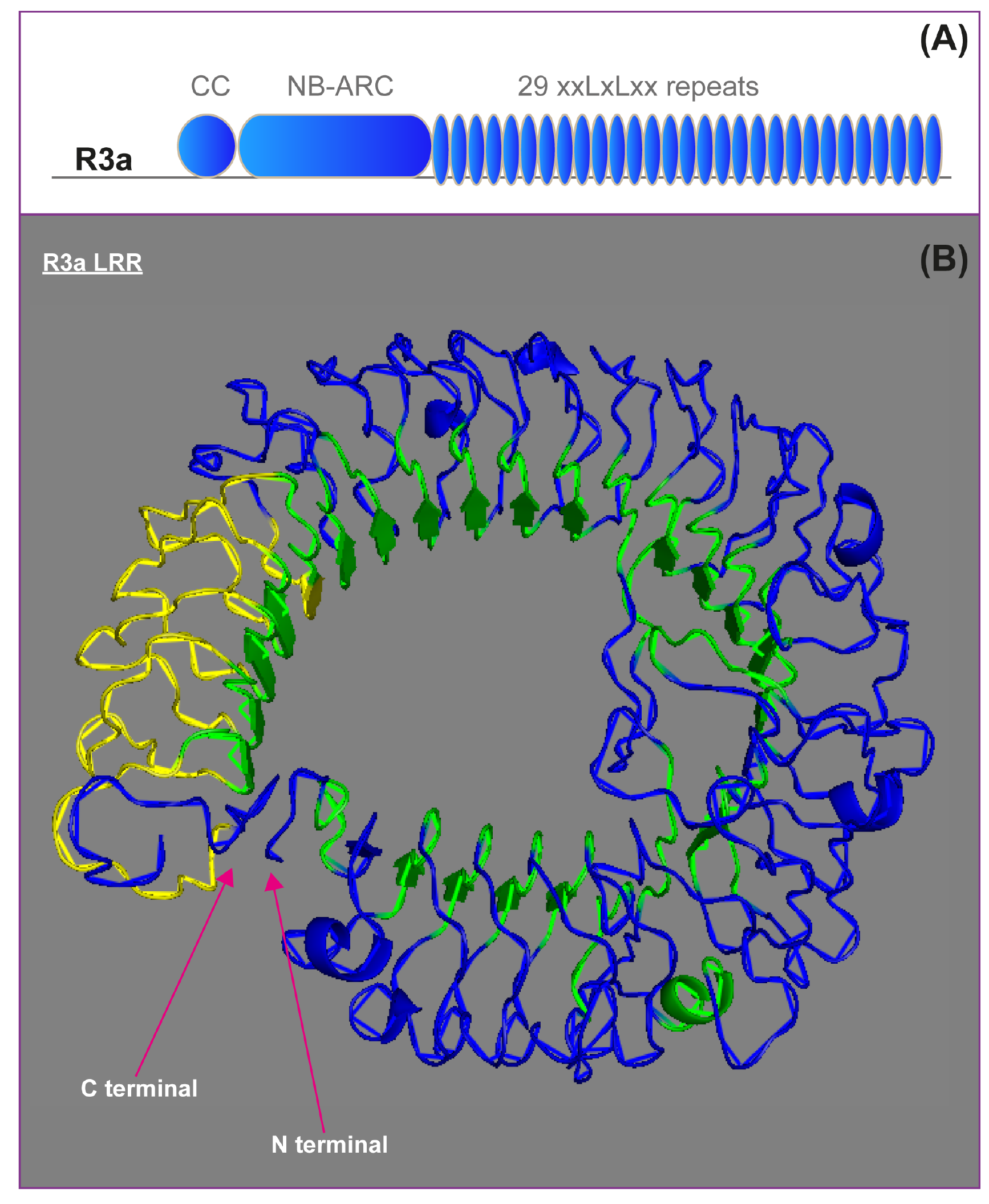
(next page). Structure homology model of the leucine rich repeat domain of R3a. A. Schematic diagram of the R3a protein. B. Structural representation of the R3a LRR domain. We generated the model by homology modeling using the IntFold server (Buenavista et al. 2012). LRRs are highlighted in green. The C-terminal region at amino acids 1148-1263 predicted by Kroj et al. (2016) to match the FAM75 domain is highlighted in yellow. The R3a protein sequence used for the analysis is GenBank accession AAW48299.1.

We conclude that R3a does not carry additional domains besides the classic CC, NB, and LRR modules of NLR proteins. Our analysis highlights the possibility of false positives among computationally predicted NLR-integrated domains. The position of predicted integrated domains needs to be checked for overlap with canonical NLR domains. Other false positives may include the DUF3542 domain, which overlaps with the CC domain of several solanaceous NLR proteins and may be an extension of the CC domain. DUF3542 was reported as the second most abundant integrated domain (Sarris et al. 2016), and was assigned to R1, another potato late blight resistance protein (Kroj et al. 2016). In R1 (GenBank accession AF447489_1), the CC domain (amino acids 406-515) overlaps with DUF3542 (amino acids 111-515) indicating that these two domains might be related.

## ACKNOWLEDGEMENTS

Research in the Kamoun Lab is funded by the Biotechnology and Biological Sciences Research Council (BBSRC), the European Research Council (ERC), and the Gatsby Charitable Foundation.

